# Development of a monoclonal antibody to a vibriophage as a proxy for *Vibrio cholerae* detection

**DOI:** 10.1101/2022.04.08.487661

**Authors:** Md. Abu Sayeed, Taylor Paisie, Meer Taifur Alam, Afsar Ali, Andrew Camilli, Jens Wrammert, Ashraful Islam Khan, Firdausi Qadri, Marco Salemi, J. Glenn Morris, Eric J. Nelson

## Abstract

Cholera is an acute watery diarrheal disease that causes high rates of morbidity and mortality without treatment. Early detection of the etiologic agent of toxigenic *Vibrio cholerae* is important to mobilize treatment and mitigate outbreaks. Monoclonal antibody (mAb) based rapid diagnostic tests (RDTs) enable early detection in settings without laboratory capacity. However, the odds of an RDT testing positive are reduced by nearly 90% when the common virulent bacteriophage ICP1 is present. We hypothesize that adding a mAb for the common, and specific, virulent bacteriophage ICP1 as a proxy for *V. cholerae* to an RDT will increase diagnostic sensitivity when virulent ICP1 phage are present. In this study, we used an *in-silico* approach to identify immunogenic ICP1 protein targets that were conserved across disparate time periods and locations. Specificity of targets to cholera patients with known ICP1 was determined, and specific targets were used to produce mAbs in a murine model. Candidate mAbs to the head protein demonstrated specificity to ICP1 by ELISA and an ICP1 phage neutralization assay. The limit of detection of the final mAb candidate for ICP1 phage particles spiked into cholera stool matrix was 8 × 10^5^ plaque forming units by Western blot analysis. This mAb will be incorporated into a RDT prototype for evaluation in a future diagnostic study to test the guiding hypothesis behind this study.

## INTRODUCTION

Cholera continues as one of the most important public health problems since 19th century, especially in resource-limited settings. Cholera can result in severe dehydration and death if untreated (1). The ongoing seventh cholera pandemic started in Indonesia in 1961 (2). Cholera remains endemic in regions of south-east Asia and Africa where there is a lack of safe drinking water, hygiene and improved sanitation (2, 3). It is estimated that 1.3 to 4.0 million cholera cases occur globally annually with 21,000 to 143,000 deaths (1, 4, 5). The frequency of cholera outbreaks is likely to rise due to globalization, rapid urbanization, and climate change (6, 7). The causative agent for cholera is toxigenic *Vibrio cholerae*, a Gram-negative facultative anaerobe. *V. cholerae* can be classified into two biotypes, classical and El Tor, more than 200 serogroups (O1-O200), and two serotypes for O1, Ogawa and Inaba. Out of all serotypes, *V. cholerae* El Tor, O1, Ogawa and Inaba are the main etiologic agents for cholera outbreaks (8, 9).

Cholera outbreaks in endemic settings follow a seasonal pattern. During outbreaks, cholera patients shed hyper-infectious *V. cholerae* as well as virulent bacteriophages (phages) (10). The proportion of cholera positive stool samples carrying virulent phage likely increases over the course of an outbreak and can reach 100% (11). It is hypothesized that the predation of virulent phages influences the seasonal pattern of cholera epidemics in cholera endemic regions (10-13). Three primary virulent phages (ICP1, ICP2, ICP3) have been found in the stool of cholera patients in Bangladesh (14, 15). ICP1, a member of *Myoviridae* bacteriophage family is the most prevalent phage excreted in cholera patient’s stool during the episode of an epidemic (14, 16, 17). ICP1 phage is specific to *V. cholerae* O1 and has been in other geographical locations including India and Africa (South Sudan and Democratic Republic of Congo (DRC)) (16-19).

According to the World Health Organization, it is estimated that more than 90% of the annual cholera cases are not reported (20). The underestimation of cholera incidence acts as a barrier for planning and implementation of acute and long-term mitigation. Lack of resources for diagnostics and appropriate surveillance system in cholera prone areas is one of the major reasons for underreporting (2, 5, 21) and delayed public health response. A rapid and accurate point of care diagnostic test can expedite cholera surveillance, response and ultimately reduce mortality and morbidity (22-24).

The gold standards for cholera diagnosis are microbial culture and polymerase chain reaction (PCR) for the detection of *V. cholerae* from stool sample. However, the sensitivity of culture method alone is approximately 70% and requires at least 2-3 days in a well-equipped microbiology laboratory with trained personnel (16, 25-28). PCR for the detection of pathogens is an alternative to the culture because of its higher sensitivity of approximately 85% (8, 29). PCR is more rapid than conventional culture, but this technique requires expensive reagents and molecular equipment as well as trained laboratory staff.

Rapid diagnostic tests (RDTs) can be used by minimally trained staff at the bed side without requiring a cold-chain or advanced equipment. More than twenty cholera RDTs have been developed (20). Most are based on immunochromatographic immunoassays, targeting *V. cholerae* O1 lipopolysaccharide antigen (30-32). Laboratory and field evaluation of RDTs showed a wide range of sensitivity and specificity of around 32 to 100% and 60 to 100%, respectively (4, 16, 24). RDT performance metrics are variable which largely limits their scope of use to cholera detection and surveillance. Our group has shown previously that virulent phage ICP1 and antibiotic exposure negatively impacts RDT performance. The odds of cholera RDT test positivity decreases by up to 90% when ICP1 phage are present (10, 16). To address this limitation, we hypothesized that adding an antibody for ICP1 to the RDT will be associated with an increase in sensitivity without compromising specificity when ICP1 phage are present in cholera stool (Fig 1). In this study, we used *in-silico, in-vitro* and *in-vivo* techniques to develop a mAb that demonstrates specificity for the ICP1 phage, with the goal to incorporate the phage mAb into the RDT and evaluate the novel RDT in a future diagnostic study.

**Fig 1.**
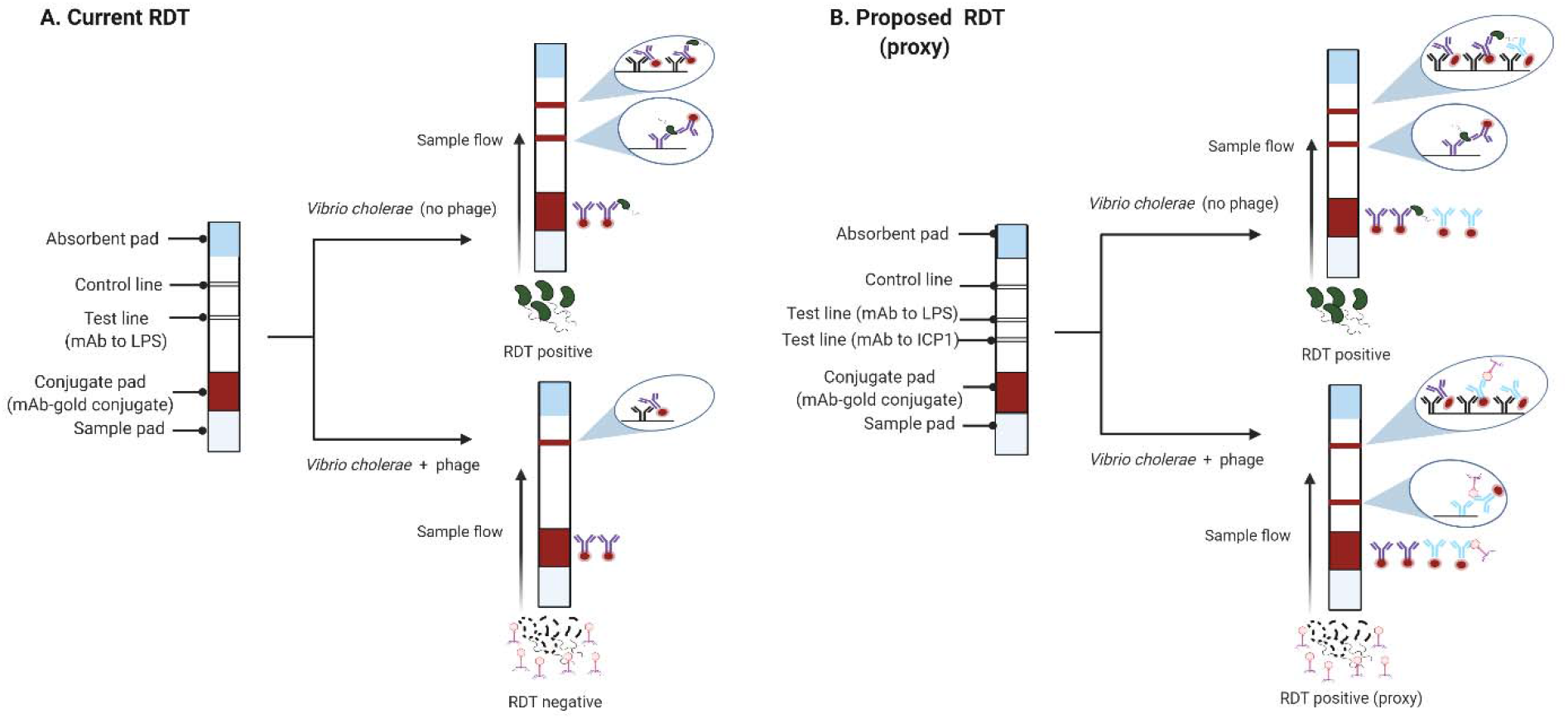
A model showing how an RDT for *V. cholerae* may fail when virulent phage (e.g. ICP1) are present in a stool sample (**A**) and how this limitation can be addressed by incorporating a mAb to phage as a proxy for *V. cholerae* when phage are present (**B**).

## METHODS

### Clinical sample collection

The sample collections analyzed were from previously published IRB approved studies; the recruitment, consent, enrollment and procedures are described (16, 19, 33, 34). In the first collection, stool samples from the Bangladesh study were obtained during September to December 2015 at a district hospital and a subdistrict hospital in the remote northern district of Netrokona (33). In the second library, stool samples from the South Sudan study were obtained during August to September 2015 at a cholera treatment center in Juba (34). The samples were collected prior to hospital administration of antibiotics; patient histories were negative for known antibiotic exposure. Lastly, microbiologic reagents were also obtained from a study in the Democratic Republic of Congo (19).

### Microbiologic procedures

#### Bacterial strains, Phage, Media, and Growth Conditions

We used the *V. cholerae* O1 strain HC1037 to isolate and prepare virulent phages ICP1, ICP2 and ICP3. This strain naturally lacks K139 prophage and is sensitive to ICP1, ICP2 and ICP3. The bacterial strain was grown at 37°C in Luria-Bertani (LB) broth with aeration or on LB agar plates (10, 14). The bacterial strain and phages used in this study are listed in Table S1. *Phage preparation, isolation, and plaque assays*. We used polyethylene glycol (PEG) precipitation method to make high titer phage stocks (15). *V. cholerae* was streaked on LB plate and incubated overnight at 37°C. A single colony from the plate was grown in LB broth to mid-exponential phase (OD_600_=0.3). Phages were added to the culture at a multiplicity of infection (MOI)=0.001 and incubated for 4-6 hours. The culture suspension was spun at 10,000 x g for 15 min at 4°C. After 0.2 μm filter sterilization of the supernatant, 0.25 volume PEG solution (20% PEG-8000; 2.5M NaCl) was added to the supernatant and incubated at 4°C for overnight for phage precipitation. Phages were pelleted by centrifugation at 10,000 x g for 25 min at 4°C. Phages were then washed with another round of PEG precipitation and finally resuspended in Phage80 buffer (0.085M NaCl, 0.1 mM MgSO_4_, 0.1 M Tris-HCl-pH 7.4). The titer of the phages was determined by plaque assay (10, 35). Phage preparation was serially diluted and incubated with mid-exponential *V. cholerae* culture for 10 min at room temperature. The mixture was added to soft LB agar (0.35% Agar) media and incubated at 37°C for 3-4 hours until plaques were observed. The number of plaques were then calculated as plaque forming unit (PFU)/ml.

### Molecular procedures

#### Cloning, expression, and purification of recombinant target proteins

The putative baseplate protein (ORF75) and head protein (ORF122) of ICP1 phage were selected to clone and express in *Escherichia coli (36-38)*. The open reading frames (ORFs) were amplified by PCR from genomic DNA. The primers were designed to include NdeI and XhoI restriction enzyme cutting sites at both ends of the amplified sequences. The PCR products and pET16b vector (Novagen) were digested with NdeI and XhoI at 37°C for 2 hours. The target sequences were cloned into pET16b by ligation using Quick Ligation™ Kit (NEB). After ligation, the recombinant plasmids were transformed into DH5 *E. coli* (Novagen) to make high copy plasmids. The cloned insert sequences were verified by colony PCR and DNA sequencing. The recombinant pET16b was then transformed into *E. coli* BL21 (Novagen) to express recombinant proteins as N-terminally His-tagged fusion proteins. A single transformed colony was picked to grow overnight at 37°C in LB broth containing 100 μg/ml ampicillin. The culture was diluted to OD_600_ 0.1-0.2 and incubated at 37°C in LB broth for 2-3 hours until the OD_600_ 0.5. Expression of recombinant proteins was induced for 4-6 hours at 37°C by adding Isopropyl β-d-1-thiogalactopyranoside to the culture at a concentration of 0.1 mM. The culture was centrifuged at 5,000 x g for 15 minutes. Before purification, an aliquot of pellet suspension and supernatant were analyzed by sodium dodecyl sulfate−polyacrylamide gel electrophoresis (SDS-PAGE) to confirm the expression of desired proteins.

The recombinant proteins were purified using His•Bind® Purification Kit (Novagen) following manufacturer’s instructions. In brief, the pellet was resuspended in Bugbuster reagent (5 ml/ gm of pellet) supplemented with Benzonase Nuclease (1 μl/ml), lysozyme (1KU/ml) and protease inhibitor (10 μl/ml). The cell suspension was incubated on a shaking platform at a slow setting for 20 minutes at room temperature (RT). After spinning at 16,000 x g for 20 minutes at 4°C, the supernatant was collected for analysis by SDS-PAGE. The pellet was resuspended and incubated again with same volume of Bugbuster reagent with lysozyme (1KU/ml) for 5 minutes at room-temperature (RT). After the addition of equal volume of 1:10 diluted Bugbuster reagent supplemented with protease inhibitor, the suspension was spun at 5,000 x g for 5 minutes. The pellet was washed with 1:10 diluted Bugbuster reagent and centrifuged at 16,000 x g for 15 minutes at 4°C. Proteins expressed as inclusion bodies were solubilized in 8 M urea. The lysate was mixed gently with 50% Ni-NTA His-bind slurry (EMD Millipore) at 4:1 ratio on a shaking platform for 60 minutes at RT. The lysate-resin mixture was carefully loaded into an empty column and washed 4 times with 8 M urea (pH-6.3). Monomeric recombinant proteins were eluted with 8 M urea (pH-5.9) and multimeric proteins were eluted with 8 M urea (pH-4.5). The purity of the proteins was further assessed by SDS-PAGE analysis and the protein concentration was measured using the Bradford method.

### Immunization and antibody production in cell culture

The purified recombinant ICP1 bacteriophage proteins were used to raise mAbs via a commercial vendor (ProMab Biotechnologies, Inc.) using a conventional hybridoma technique (28). Supernatants from 20 hybridoma clones for each of the two recombinant proteins from were received from the vendor.

### Immunologic assays of monoclonal antibody candidates

*Enzyme linked immunosorbent assay (ELISA)*. ELISA was used to screen the reactivity of ORF75 and ORF122 specific-hybridoma clones to ICP1, ICP2 and ICP3 phages (39, 40). Nunc MaxiSorp plates were coated overnight at RT with ICP1 (10^3^ PFU/well), ICP2 (10^3^ PFU/well), ICP3 (10^3^ PFU/well), formalin killed *V. cholerae* (VC;10^3^ CFU/well), ORF75 (200 ng/well), ORF122 (200 ng/well), and Bovine serum albumin (BSA; 200 ng/well). BSA and VC were used as negative controls and recombinant ORF122 or ORF75 proteins were used as positive controls. After blocking with 1% BSA-PBS, the supernatants of ORF75 and ORF122 hybridoma clones were added to the wells at 1:20 and 1:100 dilution, respectively and incubated for 1 hour at 37°C. Horseradish peroxidase-tagged goat anti-mouse IgG (Jackson ImmunoResearch) was added at 1:1000 dilution to detect antigen bound IgG mAbs. We used chromogenic substrate, 1-Step™ Ultra TMB to develop color. After stopping the reaction with 2 N H_2_SO_4_, the absorbance was measured at 450 nm using ELISA plate reader. The absorbance corresponds to the antibody binding to the coated antigens.

#### Western blot analysis

The antigens were boiled with NuPAGE SDS sample buffer containing beta-mercaptoethanol for 10 min. The wells of NuPAGE 4-12% Bis-Tris precast gel (ThermoFisher) were loaded with ICP1 (10^8^ PFU/ well), ICP2 (10^8^ PFU/ well), ICP3 (10^8^ PFU/ well), VC (5×10^5^ CFU/well), ORF122 (2 μg/well), ORF75 (2 μg/well) and BSA (2 μg/well). To determine the limit of detection (LOD), we spiked ICP1 in VC positive and ICP1 negative stool sample and prepared a 3-fold dilution series starting from 10^8^ PFU/ well. After electrophoresis at 150 V for around 40-50 mins, the proteins from unstained gel were transferred to a nitrocellulose blotting membrane using Trans-Blot turbo Transfer System (Bio-Rad). The membrane was blocked with 5% skim milk in Tris buffered saline (TBS) for overnight at 4°C. To prepare primary antibody, the supernatants of hybridoma clones were diluted in 5% skim milk-TBS-Tw (0.1%) at 1:500 dilution. The primary antibody was added on the membrane and incubated for 1 hour with gentle shaking at RT. Following washing three times with TBS-Tw (0.1%), the membrane was incubated for 1 hour with the secondary antibody, alkaline phosphatase conjugated goat anti mouse IgG (1:5000 fold diluted in 5% skim milk-TBS-Tw) with gentle shaking at RT. The membrane was then washed three times with TBS-Tw (0.1%) and developed with 5-bromo-4-chloro-3-indolyl-phosphate/nitro blue tetrazolium (BCIP/NBT) substrate for around 5 minutes. The image of protein bands was captured in a gel imager (Geldoc; Bio-Rad).

#### Phage neutralization assays

Phage neutralization assays were developed and used to test neutralization/binding by each mAb to ICP1 in a biological context (41, 42). The mAbs were diluted to 20-fold in PBS and incubated with 60-100 PFU of ICP1, ICP2 and ICP3 for 1 hour at 37°C. The phage-sample mixture was added to an exponential *V. cholerae* culture (OD_600_ 0.3) and incubated for 7-10 minutes at RT. The mixture was then added to soft LB agar (0.35% Agar) media and incubated at 37°C for 3-4 hours until plaques were observed. Phage neutralization was determined by comparing the plaque counts obtained from the assay without the mAb (only PBS).

### Statistical and bio-informatic analysis

We used VaxiJen server (VaxiJen - Drug Design and Bioinformatics Lab) and IEDB tool (National Institute of Allergy and Infectious Diseases) for predicting possible antigenic ORFs of ICP1 bacteriophage (43, 44). Clustal Omega (EMBL-EBI) was used for comparing the ORF75 and ORF122 sequences of ICP1 genomes collected from different geographical locations. In order to compare how conserved ORF75 and ORF122 were in ICP1, alignments of 29 isolates from Bangladesh and Africa (DRC) were used; analyses were at both amino acid and nucleotide levels. The data sets were used to construct a maximum likelihood (ML) phylogeny using the program IQ-TREE (45). The ML phylogeny was then used to assess temporal signal, using the program Temp-Est (46), in order to establish how conserved the ORF75 and ORF122 are in ICP1. The MSA alignment was then plotted using the R package ggmsa (http://yulab-smu.top/ggmsa/) and the temporal signal was plotted in R using ggplot2 and custom R scripts (47).

GraphPad Prism version 8 (GraphPad Software, Inc.) was used for statistical analyses and graphical presentation. The differences in antigen specific antibody responses were statistically evaluated by paired t-test. We also used the paired t-test to compare the antibody mediated phage neutralization with control. The differences were considered as statistically significant if *p* value was less than 0.05.

## RESULTS

### Selection and characterization of ICP1 protein targets for monoclonal antibody production

Eleven conserved ICP1 bacteriophage target open reading frames (ORFs) were identified and evaluated *in-silico* for immunogenic epitopes using VaxiJen and IEDB tools. The target ORFs were predicted to be highly antigenic with antigenicity scores of 0.54 to 1.03 (threshold of predicted antigen 0.4) by Vaxigen (Table S2). The targets were cross verified by IEDB to confirm antigenic epitopes (Table S2).

For further study, we selected the two putative structural proteins: a putative baseplate protein (ORF75) and a putative major head protein (ORF122). Analysis was performed on these targets to assess conservation by time and location. Conservation was found at both the nucleic acid and amino acids levels (Fig 2 A,B; Fig S2 A,B).

**Fig 2.**
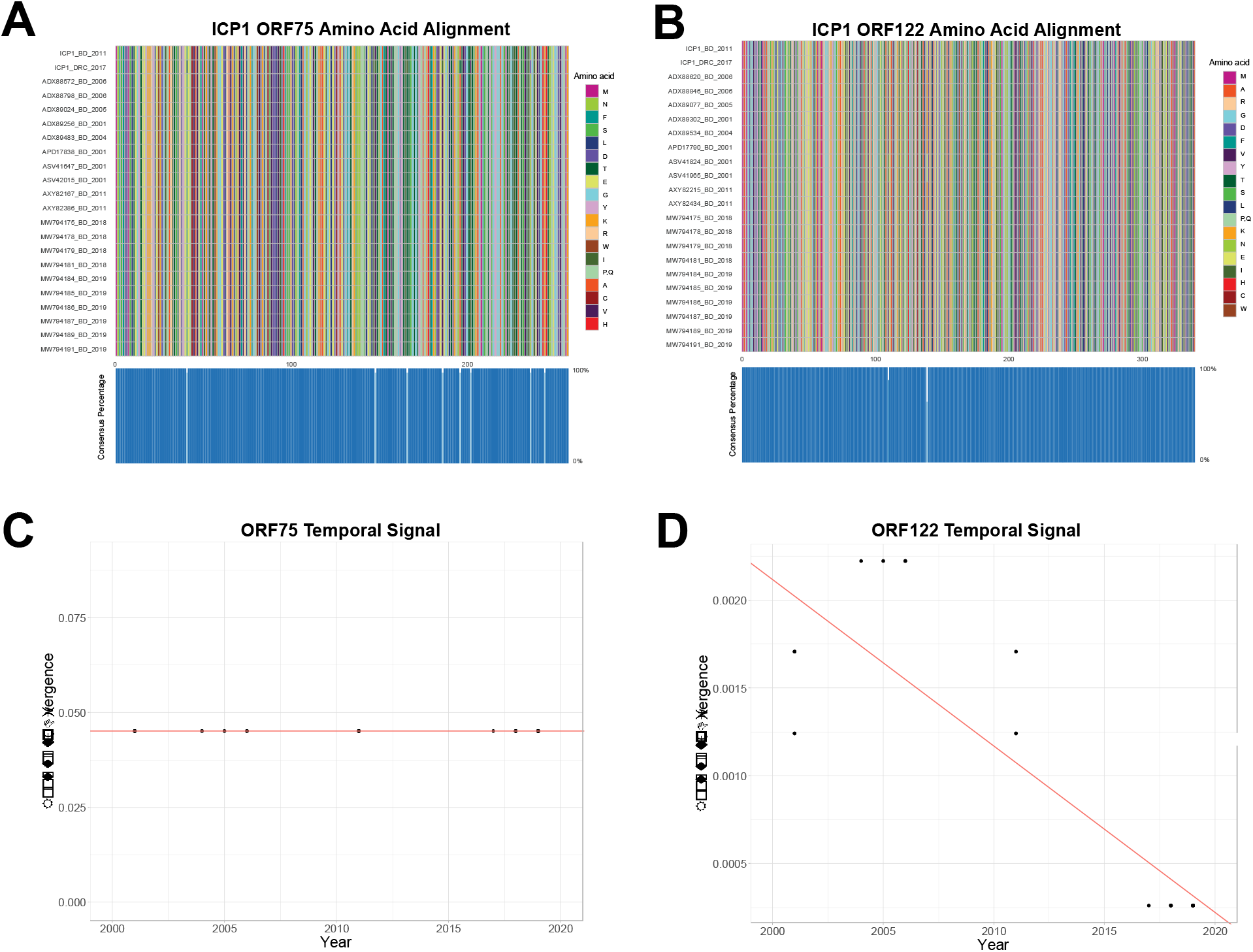
Multi-sequence alignment of ICP1 phage baseplate ORF75 **(A)** and capsular head ORF122 **(B)** amino acid sequences. Sequences from both Bangladesh (BD) and Democratic Republic of Congo (DRC). Blue boxes at the bottom of ‘A’ and ‘B’ represent the percentage of the isolates that have the same amino acid for that particular site. Temporal and divergence analysis of baseplate ORF75 **(C)** and capsular head ORF122 **(D)** nucleotide sequences from ICP1 phage isolated from Bangladesh.

However, ORF75 demonstrated higher rates of genetic diversity over time (Fig 2C, Fig S2C) compared to ORF122 (Fig 2D, Fig S2D). The target ORF75 from the type-strain ICP1 from Bangladesh (ICP1_2011_A) showed 92% (709/774) similarity at the nucleotide level and 97% (249/257) similarity at the amino acid level compared to an ICP1 isolate from Africa (DRC; ICP1_DRC_106) by a Clustal Omega sequence alignment. The target ORF122 from the type-strain ICP1 from Bangladesh (ICP1_2011_A) showed 99.8% (1018/1020) similarity at the nucleotide level and 99.7% (338/339) similarity at the amino acid level compared to an ICP1 isolate from Africa (DRC; ICP1_DRC_106); see Supplemental material Fig S1.

For Figure 2A,B displaying the amino acid multi-sequence alignments (MSA), we detected a small amount of non-synonymous (dN) mutations. We observed more dN mutations in ORF75 than in ORF122. The temporal signal in Figure 2C,D highlights the conservation of ORF75 and ORF122; the temporal signal can infer whether or not accumulating mutations are observed over time and a dataset with an accumulation of mutations overtime would be displayed with a positive slope in the temporal signal plots. A neutral slope and negative slope were observed for ORF75 (Fig. 2D) and ORF122 (Fig. 2D), respectively. Similar findings were observed at the nucleotide level (Fig. S2).

The ORF75 and ORF122 targets were screened (present/absent) by PCR in 12 phage and *V. cholerae* negative (10 from Bangladesh and 2 from Africa (South Sudan)), 2 ICP1 phage negative and *V. cholerae* positive (one from Bangladesh and one from Africa (South Sudan)), and 2 both ICP1 phage and *V. cholerae* positive stool samples (one from Bangladesh and one from Africa (South Sudan)). The ORFs were not detected in ICP1 negative stools (cholera or non-cholera). The ICP1 positive stools from both Bangladesh and Africa (South Sudan) were positive for ORF75 and ORF122 (Table S3).

### Evaluation of monoclonal antibody (mAb) candidates by ELISA

Culture supernatants of ORF75 hybridoma clones showed minimal to no reactivity to ICP1 bacteriophage in contrast to positive reactivity with purified ORF75 protein; cross-reactivity to ICP2, and ICP3 was not detected (Fig 3A). In contrast, nineteen out of twenty culture supernatants of ORF122 hybridoma clones were reactive to ICP1 and purified ORF122; cross-reactivity to ICP2 and ICP3 was not detected. The relative responses to ICP1 were significantly higher in comparison to ICP2, ICP3, formalin-killed V. cholerae whole-cell (VCWC) and BSA; but was comparable with purified ORF122 protein (Fig 3B). Given the failure of the ORF75 candidate mAbs to detect native ICP1, these candidates were eliminated from further analysis. Three ORF122 hybridoma clones including clone 5 (ICP1ORF122_mAbCL5), clone 6 (ICP1ORF122_mAbCL6) and clone 14 (ICP1ORF122_mAbCL14) were selected for further analysis based on high reactivity to ICP1.

**Fig 3.**
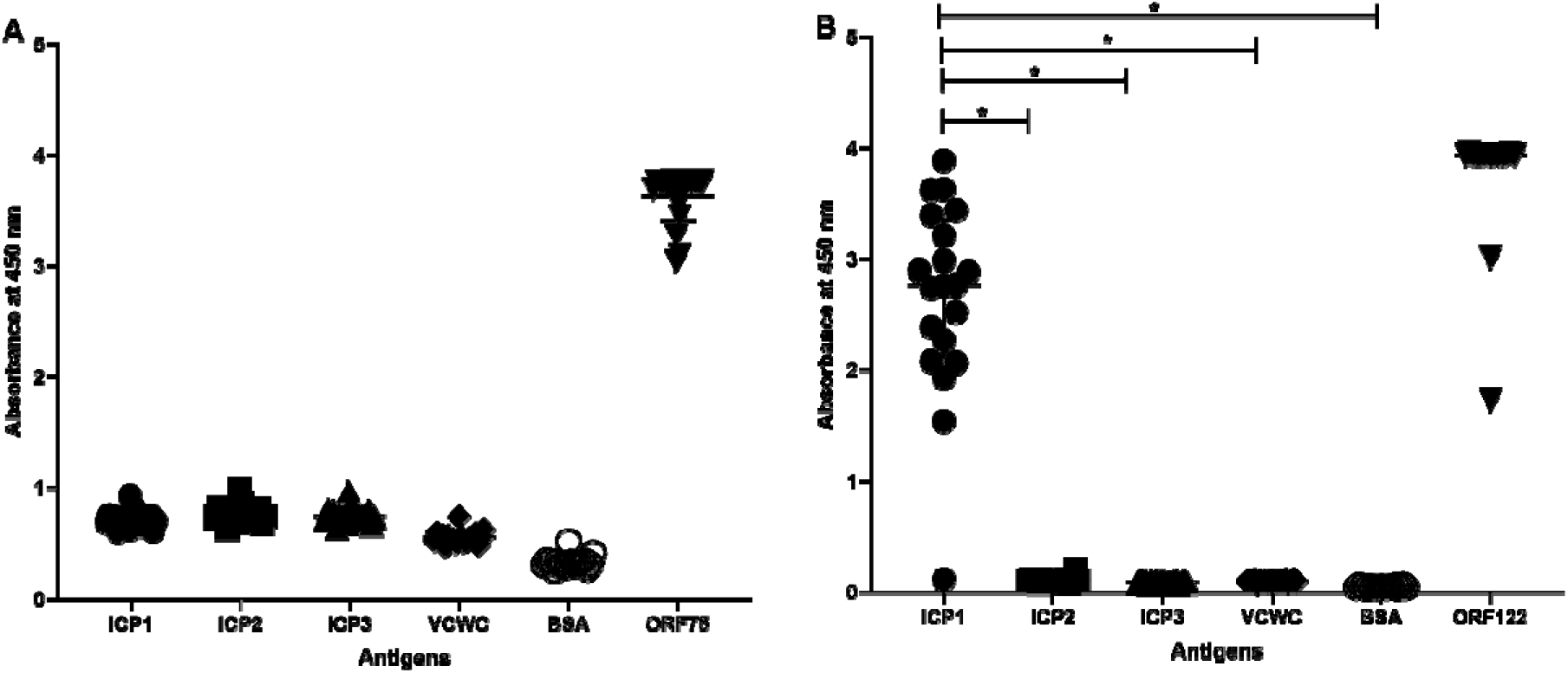
Immunoreactivity of ORF75 mAbs (n=20) (**A**) and ORF122 mAbs (n=20) (**B**) to phage particles (ICP1, ICP2, ICP3), folmalin-killed *V. cholerae* whole-cell (VCWC), bovine serum albumin (BSA) and ORF75 and ORF122 recombinant protein. Statistically significant differences (*P*<0.05) in the mean immune response from all clones are denoted with an asterisk. Symbols represents the average of three technical replicates for one mAb from one experiment; data are representative of two independent experiments.

### Evaluation of head protein monoclonal antibody (mAb) candidates by Western blot analysis

The three candidate ORF122 hybridoma clone supernatants were analyzed by Western blot. All three clones detected ICP1 as well as purified ORF122 recombinant protein. Cross-reactivity was not observed among the negative controls (ICP2, ICP3, VCWC, BSA, PBS; Fig 4A). All three candidate mAb clone supernatants detected ICP1 isolates from the disparate locations for Bangladesh and Africa (Goma DRC, Fig 4B).

**Fig 4.**
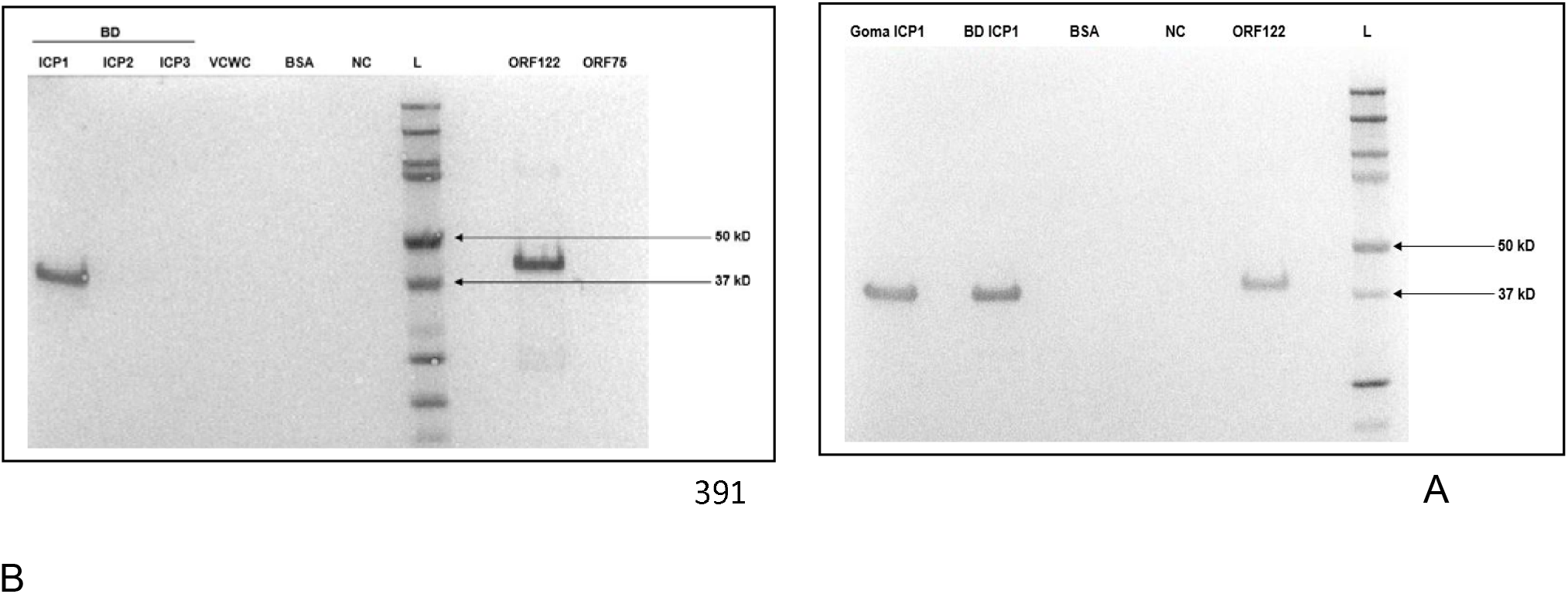
Western blot analysis of candidate ICP1ORF122_mAbCL6 against ICP1 from Bangladesh (**A**) and Goma, DRC (**B**). Negative controls are ICP2 and ICP3. BD= Bangladesh, VCWC = formalin-killed *V. cholerae* whole-cell, bovine serum albumin = BSA, NC = negative control (only PBS), L = ladder (protein marker), ORF75 and ORF122 = ICP1 recombinant proteins.

### Evaluation of head protein monoclonal antibody (mAb) candidates by phage neutralization assay analysis

We characterized the three ICP1 reactive ORF122 clone supernatants using a phage neutralization assay. All three mAb supernatant clones showed statistically significant neutralization of ICP1 (Fig 5); ICP1ORF122_mAb CL5, CL6 and CL14 were able to neutralize 31%, 42% and 39% ICP1 bacteriophage, respectively in comparison to control (only PBS). The reduction in plaque counts by phage neutralization for all the clones were statistically significant (*P*<0.001; Fig 5A).

**Fig 5.**
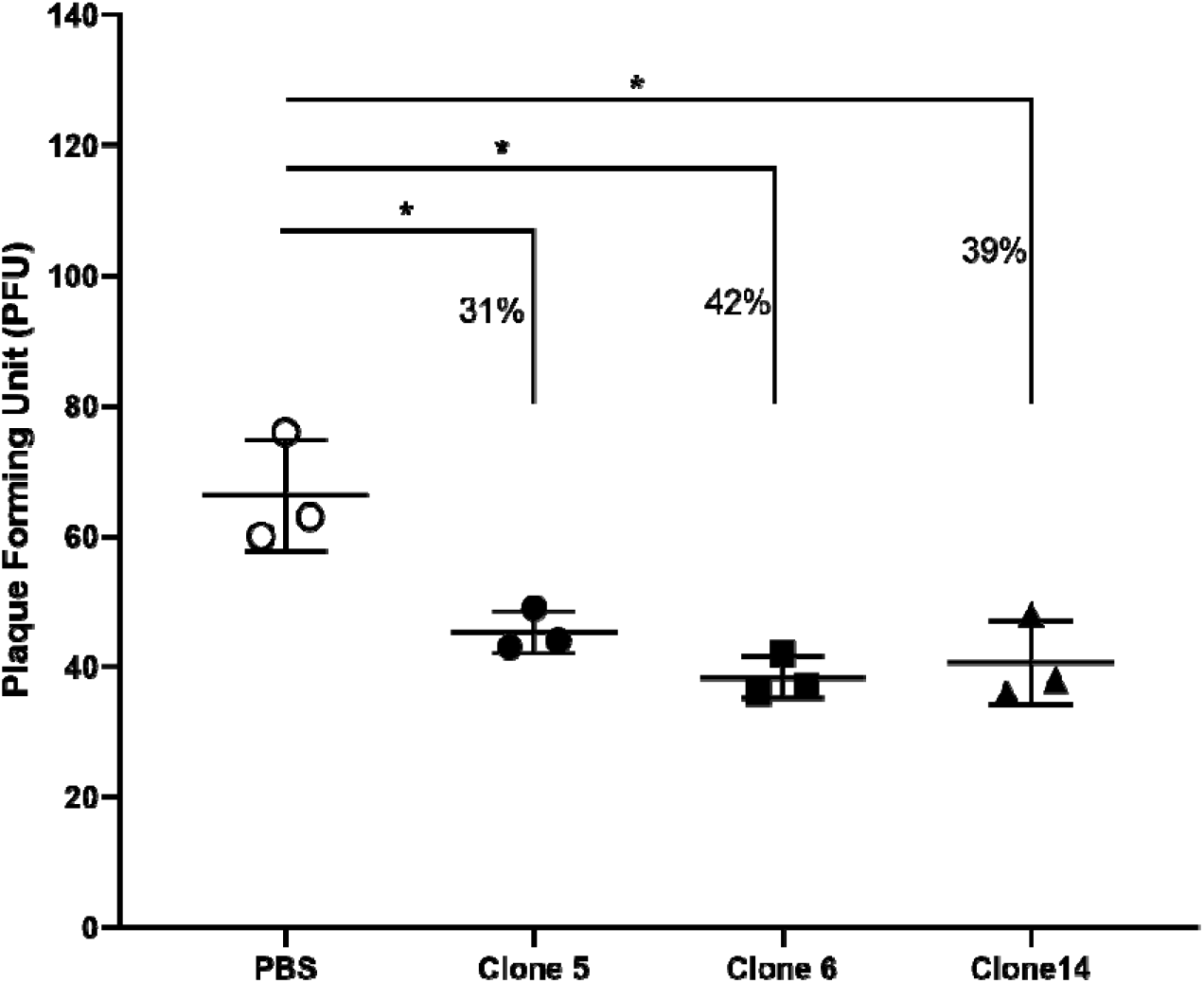
ICP1 phage neutralization by ICP1ORF122_mAb CL5, CL6 and CL14. Here, PBS= phosphate buffered saline, ICP1ORF122_mAb CL5, CL6 and CL14 represent culture supernatants from ORF122 hybridoma clones 5, 6 and 14, respectively. An asterisk denotes the statistically significant difference (*P*<0.05) in plaque counts when ORF122 mAb mediated neutralization responses are compared with the control (PBS). Each symbol represents the average of three technical replicates for one mAb from one experiment.

### Limit of monoclonal antibody detection of ICP1 bacteriophage in cholera stool matrix by Western blot analysis

We determined the limit of detection of ICP1 phage for the two final candidate clone supernatants (ICP1ORF122_mAb CL5 and CL6). We spiked ICP1 bacteriophage into ICP1 negative and *V. cholerae* negative stool samples in 3-fold dilution series. We found that both CL5 and CL6 culture supernatants (1:500) were able to detect down to 8 × 10^5^ PFU of ICP1 bacteriophage by Western blot (Fig 6).

**Fig 6.**
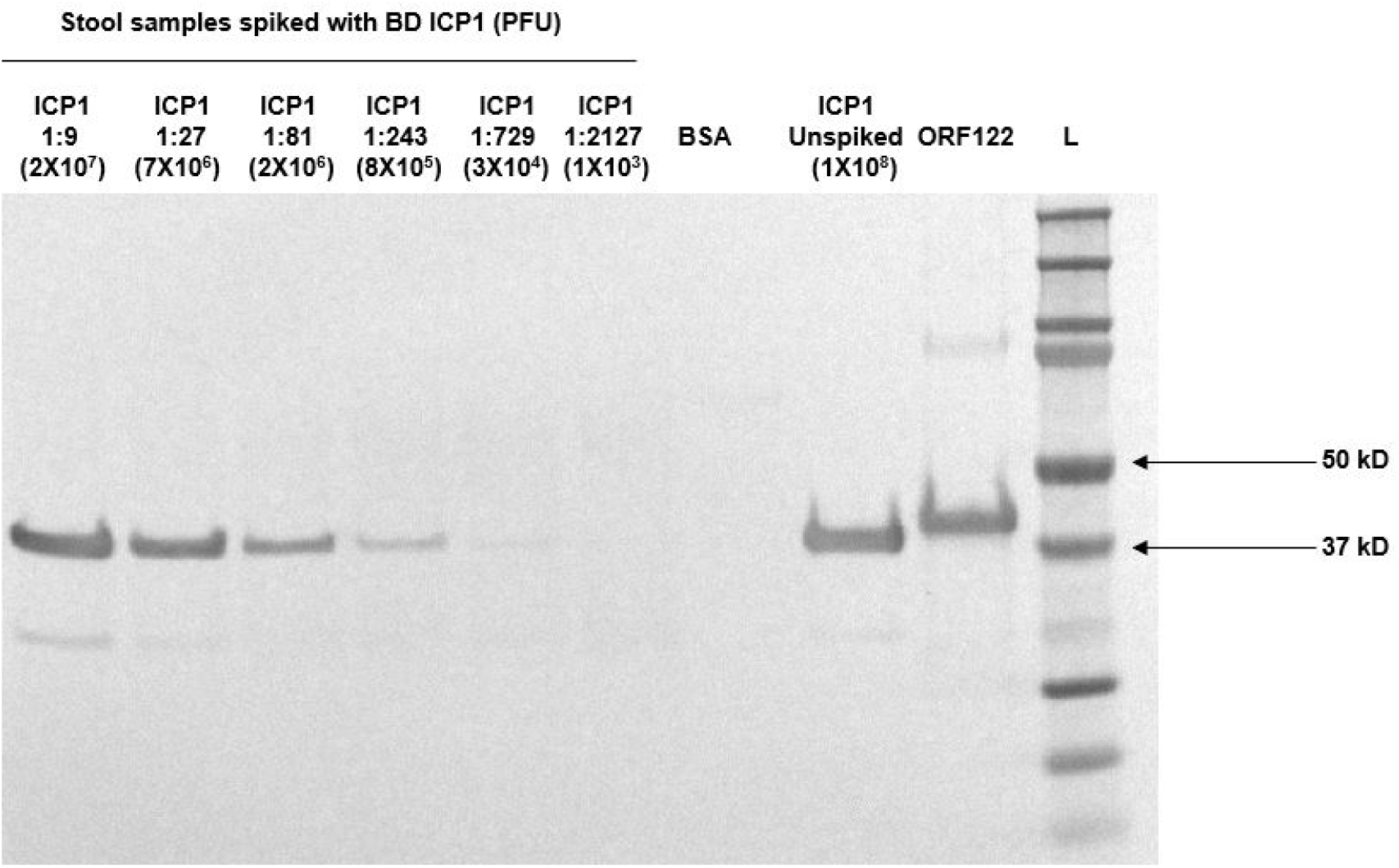
Determination of the limit of detection (LOD) of ICP1ORF122_mAbCL6 against ICP1 in cholera stool supernatant. ICP1 was serially diluted in cholera stool known to be vibriophage negative (ICP1, ICP2, and ICP3 negative) in 3-fold dilution series. The concentration of the neat ICP1 stock was 2 × 10^10^ PFU/ml. The lane with 1:243 dilution represents 8 × 10^5^ PFU of ICP1 phage. Here, bovine serum albumin=BSA, ORF122= ICP1 recombinant protein and L= ladder (protein marker).

## DISCUSSION

In this study, we aimed to develop a mAb against the common virulent vibriophage ICP1 as a critical step towards addressing limitations with current cholera RDTs. Our guiding hypothesis is that adding a mAb for ICP1 to the existing RDT as a proxy for *V. cholerae* will increase sensitivity when ICP1 degrades the primary *V. cholerae* target. We used an *in-silico* approach to identify immunogenic protein targets that were conserved and specific to cholera patients. Candidate proteins were expressed for mAb production, and mAbs to the head protein (ORF122) demonstrated specificity to ICP1 by both ELISA and a phage neutralization assay. The mAb to the head protein (ORF122) was able to detect ICP1 at biologically meaningful concentrations by Western blot analysis when ICP1 was spiked into cholera stool matrix. This mAb will be incorporated into an RDT prototype for evaluation in a clinical study to test our guiding hypothesis.

This approach is innovative in that we sought to develop a mAb to a pathogen-specific phage as a proxy for the bacterial pathogen. However, the durability of the approach is vulnerable if the antigenicity of the epitope varies across time and place. The strong selective pressures between bacterial ‘prey’ and bacteriophage ‘predator’ drive elaborate mechanisms of phage immunity and escape, and ultimately genetic diversity. That said, genes for the candidate ICP1 structural proteins were found to be conserved. In prior analyses, the baseplate protein (ORF75) was conserved at near 100% similarity and the head protein (ORF122) was conserved at more than 99% similarity at both amino acid and nucleic acid levels (17). With additional data, we found the baseplate gene for ORF75 was more divergent compared to ORF 122 across time and location. Both proteins are unlikely to be present in non-cholera patients given that the ORFs were not detected by PCR in non-cholera patient diarrheal stool, and cross-reactivity between phages is unlikely given that no significance sequence homology beyond ICP1 was identified, including within *Myoviridae*.

The other vulnerability of our approach is that the mAb might have cross-reactivity or degrade in cholera stool matrix which harbors proteases (48, 49). Monoclonal antibodies were raised to recombinant ORF75 and ORF122 proteins, however the mAb to the baseplate protein (ORF75) failed to bind native ICP1 by ELISA, Western blot and phage neutralization assays. This failure might be due to the lesser abundance of the epitope in the native ICP1 phage particle, post-transcriptional modification or possibly epitope masking. This is consistent with a previous study showing that the staphylococcal phage major capsid protein was highly immunogenic, whereas the baseplate protein was found to be non-immunogenic in mice (50). On the other hand, the supernatants of the clones raised with the capsid protein ORF122 were reactive to the native ICP1 phage particle by ELISA, Western blot and phage neutralization assays. With respect to RDT development, the candidate mAbs were able to detect ICP1 alone, without cross-reactivity to ICP2 or ICP3. During western blot analysis, the cholera stool matrix with a ICP1 spike-in did not detectably interfere with ICP1 detection by the candidate mAbs.

These findings should be viewed within the context of the study limitations. First, the mAb did not fully ablate ICP1 in the viral neutralization assay. The mAb to ORF122 reduced plaque formation by 30-40% ICP1 which was less than expected given its specificity. While this modest result is consistent with a similar study on anti-T4 head antibodies neutralizing T4 phage activity in *E. coli* (51), further optimization of the assay may be needed. Alternatively, the modest neutralization may be the result of cross-linking at the capsular head of ICP1 phage particles, which may leave the apparatus for binding and injection into the bacterial host operative (51, 52). Second, the scope of investigation of the mAb cross-reactivity was limited and will be improved upon by prototyping the RDT and a prospective diagnostic study in cholera and non-cholera patients. Third, we tested ICP1 spiked in cholera stool matrix alone and we did not have access to *V. cholerae* positive stool samples with or without ICP1 phage at native concentrations. While the limit of detection of ICP1 was lower than that anticipated in cholera stool, data are limited on the native concentrations of ICP1 across the time course of disease. Lastly, the exact epitopes that the mAbs bind remain unmapped.

Despite limitations, our work has significant implications. The mAb to the head protein (ORF122) developed herein can be used for making an enhanced RDT to detect ICP1 as a proxy for *V. cholerae*. In a prospective diagnostic study, we will evaluate the performance of the enhanced RDT across the course of cholera outbreaks given that cholera patients are more likely to shed virulent phage at the latter outbreak periods (11, 12, 16).

## Acknowledgements

We thank the patients for participating in the studies from which the clinical samples were obtained. We are grateful to Randy Autrey and Krista Berquist for their administrative expertise, as well as the UF Emerging Pathogens Institute and the UF Department of Pediatrics for providing vital infrastructure.

## Data availability

Data analyzed are presented within the manuscript and online supplementary material.

## Financial Support

This work was supported by a grant from the Wellcome Trust to EJN [DP5OD019893] and internal support from the Emerging Pathogens Institute, and the Departments of Pediatrics and the Department of Environmental and Global Health at the University of Florida, and by a grant from the NIH (USA) to AC [AI055058].

## Disclaimer

The funders had no role in the study design, data collection and analysis, decision to publish, or preparation of the manuscript.

## Potential conflicts of interest

All authors: No reported conflicts.

## SUPPLEMENTARY MATERIALS

**Fig S1.**
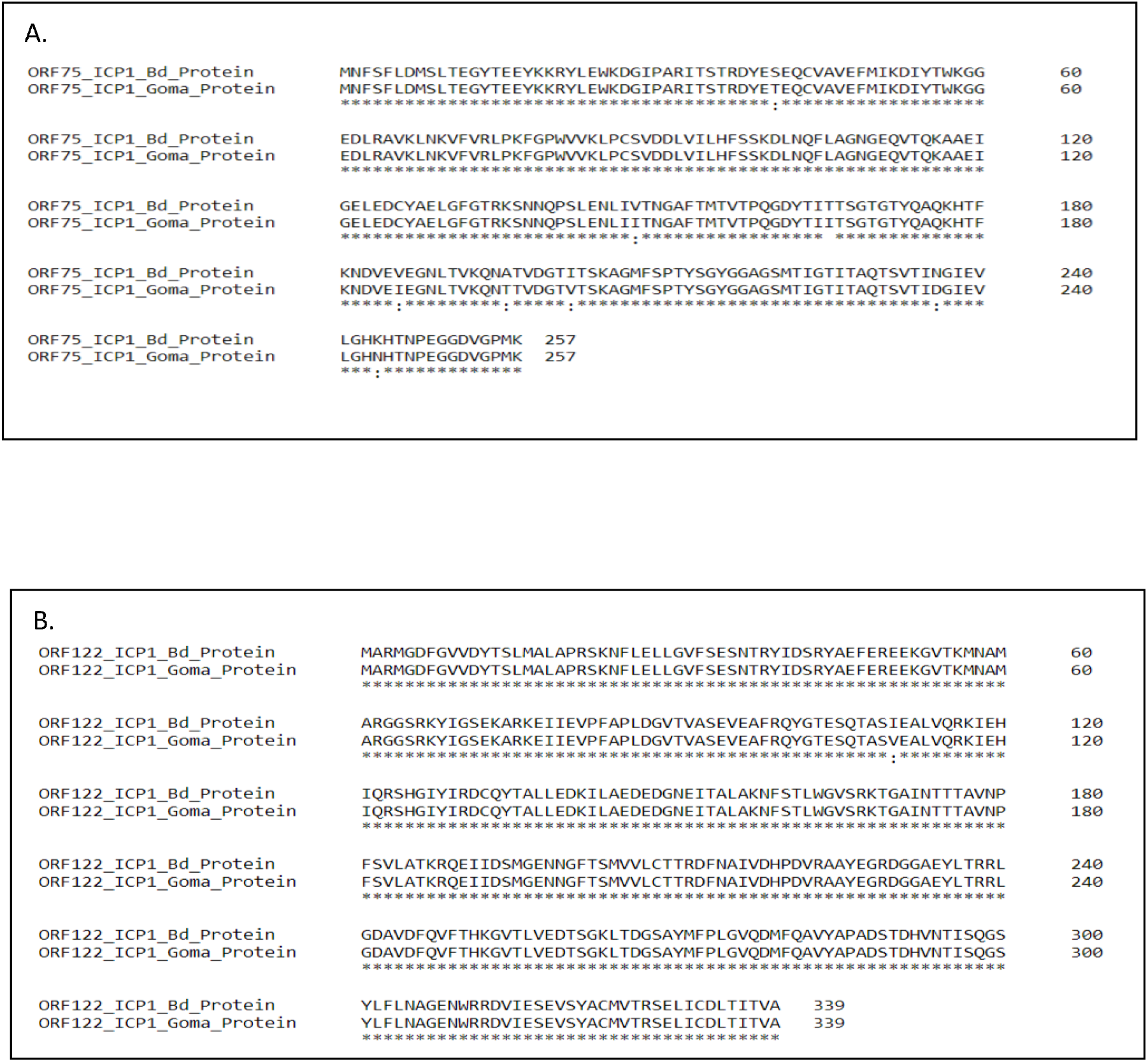
Alignment of Bangladesh (Bd) (14) and Goma ICP1 (19) bacteriophage ORF75 (**A**) and ORF122 (**B**) protein sequences by Clustal Omega (EMBL-EBI). Alignment is shown by symbols. The asterisk symbol represents identical amino acid, colon represents similar amino acid and any gaps in the alignment represent mismatched amino acid.

**Fig S2.**
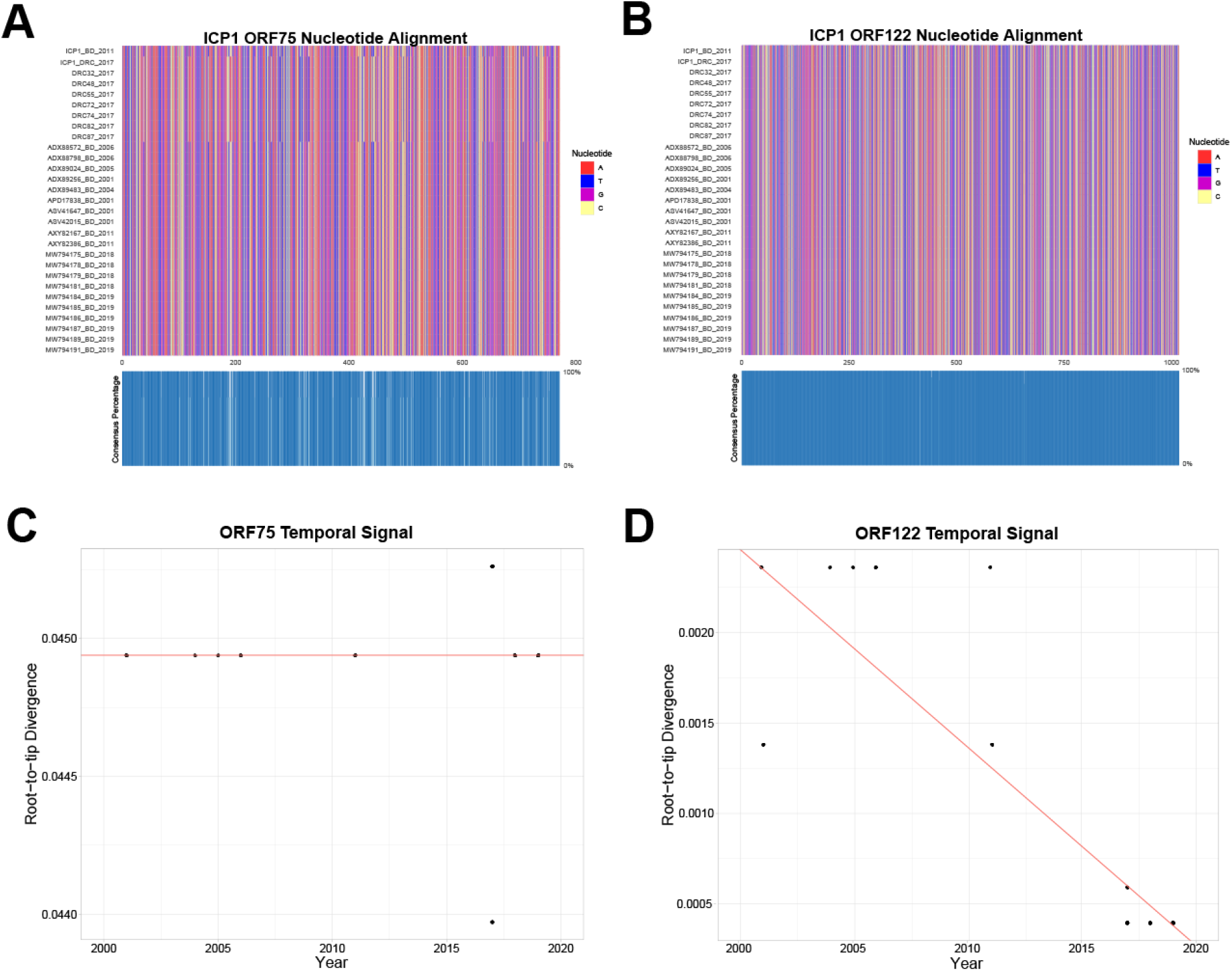
Multi-sequence alignment of ICP1 phage baseplate ORF75 **(A)** and capsular head ORF122 **(B)** nucleotide sequences. Sequences from both Bangladesh (BD) and Democratic Republic of Congo (DRC). Blue boxes at the bottom of A and B represent the percentage of the isolates that have the same nucleotide for that particular site. Temporal and divergence analysis of baseplate ORF75 (**C**) and capsular head ORF122 (**D**) nucleotide sequences from ICP1 strains isolated in Bangladesh.

**Table S1.** Microbiologic and molecular reagents

| **Reagent** |  |  |  |
| --- | --- | --- | --- |
| **Bacteria** | **Strain** | **Description** | **Reference** |
| *V. cholerae* | HC1037 | O1 Ogawa serogroup, isolated from Haiti, SmR | (53) |
| **Bacteriophage** | **Strain (host)** | **Description** | **Reference** |
| ICP1 | ICP1_2011_A<br>(*V. cholerae* O1) | *Myoviridae*, isolated from Bangladesh | (14) |
| ICP1 | ICP1_DRC_106<br>(*V. cholerae* O1) | *Myoviridae*, isolated from Goma, DRC | (19) |
| ICP2 | ICP2_2004_A<br>(*V. cholerae* O1<br>and non-O1) | *Podoviridae*, isolated from Bangladesh | (14) |
| ICP3 | ICP3_2007_A<br>(*V. cholerae* O1<br>and non-O1) | *Podoviridae*, isolated from Bangladesh | (14) |

**Table S2.** List of immunogenic ICP1 core ORFs.

| Predicted Protein | Function | Predicted Antigenicity (rank)<sup>a</sup> | Length (nt) | No of predicted epitopes<sup>b</sup> | Length (aa) |
| --- | --- | --- | --- | --- | --- |
| ORF66 | hypothetical protein | 1.03 (1) | 228 | 3 | 75 |
| ORF154 | hypothetical protein | 0.92 (2) | 132 | 1 | 43 |
| ORF158 | hypothetical protein | 0.83 (3) | 258 | 3 | 85 |
| ORF9 | hypothetical protein | 0.81 (4) | 159 | 2 | 52 |
| ORF206 | hypothetical protein | 0.77 (5) | 219 | 3 | 72 |
| ORF214 | hypothetical protein | 0.75 (6) | 189 | 3 | 62 |
| ORF75 | putative baseplate assembly protein | 0.73 (7) | 774 | 7 | 257 |
| ORF1 | hypothetical protein | 0.73 (8) | 108 | 1 | 35 |
| ORF195 | hypothetical protein | 0.71 (9) | 162 | 2 | 53 |
| ORF52 | hypothetical protein | 0.67 (10) | 183 | 4 | 60 |
| ORF122<sup>c</sup> | putative major head protein | 0.54 (17) | 1020 | 14 | 339 |

<sup>a</sup> Predicted antigenicity score of initially selected 11 ORFs by Vaxijen v2. The threshold for this model is 0.4 for a probable antigen. The rank is based on the antigenic score among 50 core ORFs (49 conserved core and 1 divergent core (c)) of ICP1 bacteriophage (17).
<sup>b</sup> Number of predicted B-cell epitopes by IEDB analysis. This analysis was done using Bepipred Linear Epitope Prediction 2.0 model.

**Table S3.** Molecular screening for predicted immunogenic targets in cholera and non-cholera stools.

| **Bangladesh samples** | **ID** | **ORF75<sup>a</sup>** | **ORF122<sup>b</sup>** |
| --- | --- | --- | --- |
| *V. cholerae* negative samples (n=10) |  |  |  |
| S1 | RN22 | - | - |
| S2 | RN23 | - | - |
| S3 | RN24 | - | - |
| S4 | RN25 | - | - |
| S5 | RN26 | - | - |
| S6 | RN27 | - | - |
| S7 | RN28 | - | - |
| S8 | RN29 | - | - |
| S9 | RN30 | - | - |
| S10 | RN31 | - | - |
| *V. cholerae* positive samples (n=2) |  |  |  |
| S1 (ICP1-) | RN3 | - | - |
| S2 (ICP1+) | VCP 12 | + | + |
| **South Sudan samples** |  |  |  |
| Random *V. cholerae* negative sample (n=2) |  |  |  |
| S1 | 1001 | - | - |
| S2 | 1002 | - | - |
| *V. cholerae* positive sample (n=2) |  |  |  |
| S1 (ICP1-) | 1008 | - | - |
| S2 (ICP1+) | 1086 | + | + |

<sup>a</sup> represents the PCR amplification results for the target ORF75, a putative baseplate protein
<sup>b</sup> represents the PCR amplification results for the target ORF122, a putative ICP1 bacteriophage head protein.

## Notes

### Competing Interest Statement

The authors have declared no competing interest.

